# miR-100-5p downregulates mTOR to suppress the proliferation, migration and invasion of prostate cancer cells

**DOI:** 10.1101/2020.07.30.228676

**Authors:** Su-Liang Li, Yun Ye, Jian-Jun Wang

## Abstract

**Background:** Previous studies have shown that miR-100-5p expression is abnormal in prostate cancer. However, the role and regulatory mechanism of miR-100-5p requires further investigation. Thus, the aim of this study was to observe the effects of miR-100-5p on the proliferation, migration and invasion of prostate cancer (PCa) cells and to explore the potential related regulatory mechanism.

**Methods:** Differential miRNA expression analysis was performed using next-generation sequencing (NGS) in the PCa cell line LNCaP and the normal prostatic epithelial cell line RWPE-1. The expression levels of miR-100-5pwere detected using real-time fluorescence quantitative PCR (qRT-PCR). LNCaP cells were transfected with NC-mimics or miR-100-5p mimics by using liposome transfection. Moreover, the CCK-8 proliferation assay, cell scratch assay and Transwell assay were used to detect the effects of miR-100-5p on cell proliferation, migration and invasion. In addition, the potential target gene of miR-100-5p was predicted, and the influence of miR-100-5p on the expression of mTOR mRNA by qRT-PCR and the expression of mTOR protein was detected by western blot and immunohistochemical staining.

**Results:** Differential expression analysis of high-throughput sequencing data showed low expression of miR-100-5p in the PCa cell line LNCaP. It was further confirmed by qRT-PCR that the expression of miR-100-5p in LNCaP cells was significantly lower than that in RWPE-1 cells (*P*<0.01). miR-100-5p expression in LNCaP cells was markedly upregulated after transfection with miR-100-5p mimics (*P*<0.01), while cell proliferation, migration and invasion capacities were clearly reduced (*P*<0.01), and mTOR mRNA and protein expression was also substantially lowered (*P*<0.01). Finally, we further confirmed by immunohistochemical staining that miR-100-5p regulated the expression of mTOR.

**Conclusion:** miR-100-5p is expressed at low levels in LNCaP cells, and it can suppress LNCaP cell proliferation, migration and invasion, the mechanism of which is related to downregulating the expression of mTOR.

## Introduction

PCa is the most common malignant tumor of the male reproductive system. According to the latest statistics, a projected 191,930 new cases of PCa will be diagnosed, accounting for more than 1 in 5 new diagnoses of male tumors. An estimated 33,330 men may die of the disease in the USA in 2020, and mortality due to PCa accounts for 10% of all cancer deaths.^1^ With an increasingly aging population and changes in diet, the incidence of prostate cancer in Asia is increasing year by year.^2^ MicroRNAs (miRNAs) are highly conserved noncoding single-stranded RNAs consisting of 21-24 nucleotides that regulate the expression of target genes through complete or incomplete complementarity binding with target genes and play an important role in the gene regulatory network.^3,4^ Recent studies have found that miRNAs, as oncogenes or tumor suppressor genes, play an important role in the development and progression of malignant tumors.^5^ A large number of studies have confirmed that miRNA expression is dysregulated in the occurrence and development of prostate cancer, and there is differential expression between prostate cancer patients and the normal population.^6,7^ MiR-100-5p is an important member of the miRNA family, located on chromosome 11q24.1 and highly conserved. MiR-100-5p has abnormal expression in many malignant tumors, including prostate cancer and is involved in biological behaviors such as proliferation, migration and invasion of tumor cells, but the mechanism of action is still unclear.^8^ In this study, we observed changes in the expression of miR-100-5p in the PCa cell line LNCaP and its effect on cell proliferation, migration and invasion ability and explored its potential molecular mechanism.

## Methods

### Cell culture

LNCaP (PCa cell line) and RWPE-1 (normal prostatic epithelial cells) were purchased from American Type Culture Collection (Manassas, VA, USA). LNCaP cells and RWPE-1 cells were grown in Roswell Park Memorial Institute (RPMI)-1640 (Gibco; Thermo Fisher Scientific, Inc., Waltham, MA, USA). All culture media were supplemented with 10% fetal bovine serum (Gibco; Thermo Fisher Scientific), 100 U/ml penicillin and 100 mg/ml streptomycin (HyClone; GE Healthcare Life Sciences). All cells were routinely cultured in a humidity incubator at 37°C and 5% CO2, and logarithmic-phase cells with good growth conditions were selected for subsequent experiments.

### Construction of the cDNA library

MiRNAs were isolated and purified by gel electrophoresis, and cDNA was synthesized by reverse transcription. The miRNA sequencing library was obtained by PCR amplification. cDNA libraries were tested using an Illumina Hi Seq 2500 sequencer (GUANGZHOU RIBOBIO CO., LTD).

### Differential expression analysis of miRNA

Edger analysis was used to analyze the significance of differences in miRNA expression in each group and calculate the P value, for which the -log10p value was calculated and the differentially expressed miRNAs were screened.

### RNA preparation and RT-PCR

We isolated total RNA and total miRNA from cells by using TRIzol (Life Technologies) reagent and the mirVana miRNA Isolation Kit (Ambion). We treated total RNA with TURBO DNase (Ambion) after purification and used a high-capacity cDNA reverse transcription kit (Takara) to reverse transcribe RNA into first-strand cDNA. We purchased primers against mRNAs from Shanghai Sangon Biotech Co., Ltd. Each experiment was carried out in triplicate, and the mean value of the three-cycle threshold was used for further analysis. Cel-miR-39 was used as an internal control. The expression values of miRNAs were normalized to the U6 values, and the relative quantification analysis was performed using the 2^-ΔΔCt^ method. The primers for miR-100-5p were F: 5’-GAACCCGTAGATCCGAACT-3’ and R: 5’-CAGTGCGTGTCGT GGAGT-3’. The primers for U6 were F: 5’-CTCGCTTCGGCAGCACA-3’ and R: 5’-AACG CTTCACGAATTTGCGT-3’. The primers for mTOR were F: 5’-CTGGGACTCAAATG TGTGCAGTTC-3’ and R: 5’-GAACAATAGGGTGAATGATCCGGG-3’. The primers for β-actin were F: 5’-TCCTCCCTGGAGAAGAGCTA-3’ and R: 5’-TCAGGAGGAGCAATGATCTTG-3’. The amplification conditions were 95°C for 10 min, followed by 40 amplification cycles of 95°C for 10 s and 60°C for 30 s. The ABI Prism 7500 Sequence Detection System (Applied Biosystems) was used to perform real-time PCR.

### Cell transfection

miR-100-5p mimics and the corresponding control (miR-NC) were synthesized by RIBOBIO Co., Ltd. (Guangzhou, China) and transfected into LNCaP cells by Lipofectamine 2000 (Invitrogen, New York, USA) according to the manufacturer’s specifications. The results were examined by qRT-PCR, and further experiments were also carried out after transfection for 48 h.

### Cell proliferation assay

The cells were proportionally diluted with culture medium and the OD value was measured after adding CCK-8 reagent for 8 h to generate a standard curve. The cell suspension (5000 cells/100 µl/well) was inoculated in a 96-well plate, and 10 µl CCK-8 solution was added to each well. The sample was incubated for 4 h, and the OD value was measured at 450 nm.

### Cell scratch assay

LNCaP cells (5 × 103 cells/well) were cultured in a 24-well plate in RPMI-1640 culture medium with 10% FBS. The culture plate was incubated overnight at 37°C in a humidified CO2 incubator. Next, the culture medium was removed, and the adherent cell layer was scratched with a sterile 200 μL pipette tip. We washed away the cell fragments with phosphate-buffered saline (PBS). Images of the scratch area at 0 h and 24 h were taken using a built-in camera in the microscope (40x magnification). Data were evaluated using TScratch imaging software (CSE Lab., ETH, Zurich) to calculate the percent wound area.^9^

### Transwell assay

The Transwell chamber was treated with Matrigel (Solarbio, Beijing, China). After transfection, 2×10^5^ LNCaP cells and NC control cells were cultured in the upper chamber (Solarbio, Beijing, China) with serum-free medium. RPMI-1640 medium containing 10% FBS was added to the lower chamber, and the Transwell chamber was removed after continued culture for 24 h and maintained at 37°C in 5% CO2 overnight. Subsequently, cells were fixed with methanol for 10 min, stained with crystal violet (Yuanye, Shanghai, China) and then analyzed by microscopy. The number of cells that had crossed the membrane was calculated, and the experiment was repeated three times.

### Western blotting

The cell harvesting and extraction of proteins followed the method described previously.^10^ A protein assay kit (Bio-Rad, Hercules, CA, USA) was used to determine protein concentrations, and SDS-PAGE was used to separate the samples according to molecular weight. The blotted membrane was incubated with rabbit monoclonal anti-mTOR (1:1,000; ab32028; Abcam), β-actin (1:1,000; ab8227; Abcam) which was used as a loading control and horseradish peroxidase (HRP)-coupled goat anti-rabbit IgG H&L (1:5,000; ab97051; Abcam) as a secondary antibody. Primary antibody dilutions were incubated with the membranes overnight at 4°C.

### Tumorigenesis studies of animals

All procedures involving mice were carried out under protocols approved by Committees of Animal Ethics and Experimental Safety of The First Affiliated Hospital of Xi’an Medical University (protocol number xyfy2019-192) and in accordance with National Health and Medical Research Council guidelines. The methods were performed in accordance with the approved guidelines by Xi’an Medical University. Twenty six-week-old male nude mice (Air Force Medical University, China) were housed in a Specific Pathogen Free barrier facility provided with HEPA filtered air with free access to sterile, standard rodent chow, and sterile water. Environmental enrichment for the animals was provided with tissue paper and cardboard. Mice were randomly divided into two groups and administered by intraosseous injected with 2.0×10^6^ tumor cells in 50% Matrigel™ (Falcon, NJ, USA), LNCaP cells transfected with miR-100-5p mimics (A group) and only LNCaP cells (B group, control). Anaesthesia was by intraperitoneal ketamine 100 mg/kg and xylazine 10 mg/kg injection.Pathological staining and immunohistochemical staining were performed after two months. Animals were monitored daily to ensure they were not experiencing distress. Humane endpoints based on body weight loss,body condition scoring and metastatic progression were in place. Mice were euthanized with CO2. No unexplained mortality occurred in these studies.

### Immunohistochemical analysis

Paraffin-embedded tissues were dewaxed, hydrated with ethanol, and incubated with 0.3% H_2_O_2_ to eliminate endogenous peroxidase activity. The primary antibody was added and incubated at 4°C overnight, and the secondary antibody was incubated at room temperature for 1 h. The immunohistochemical results were observed by a Nikon Eclipse 80i microscope, and positive results for brown granules were mainly distributed in the cytoplasm. Positive expression was quantitatively analyzed by Image-ProPlus 5.0 analysis software, and the integral optical density (IOD) value of positive staining in each visual field was calculated. The mean value of IOD of 5 visual fields was taken as the final value of each group.

### Prediction of targeted relationship

MiRGator 3.0 software and TargetScan Human 7.1 were used to predict the targeting relationship between miR-100-5p and mTOR.

### Statistical analyses

SPSS 20.0 software (SPSS Inc., Chicago, IL, USA) for Windows was used to perform all statistical analyses. The data are presented as the mean ± SD. Nonparametric data were analyzed by 2-tailed Mann-Whitney U-tests. P<0.05 was selected to indicate a statistically significant difference.

## Results

### Differential expression analysis of miRNAs

Compared with those in the normal prostate epithelial cells RWPE-1, 29 miRNAs in prostate cancer cells LNCaP were significantly differentially expressed (figure 1). Notably, miR-100-5p, miR-584-5p and miR-125b-1-3p were downregulated in LNCaP cells, whereas miR-375, miR-200c-3p and miR-141-3p expression levels were upregulated (figure 2). The qRT-PCR results further confirmed that the expression of miR-100-5p in LNCaP cells was significantly lower than that in RWPE-1 cells (P<0.01) (figure 3).

**Fig 1.**
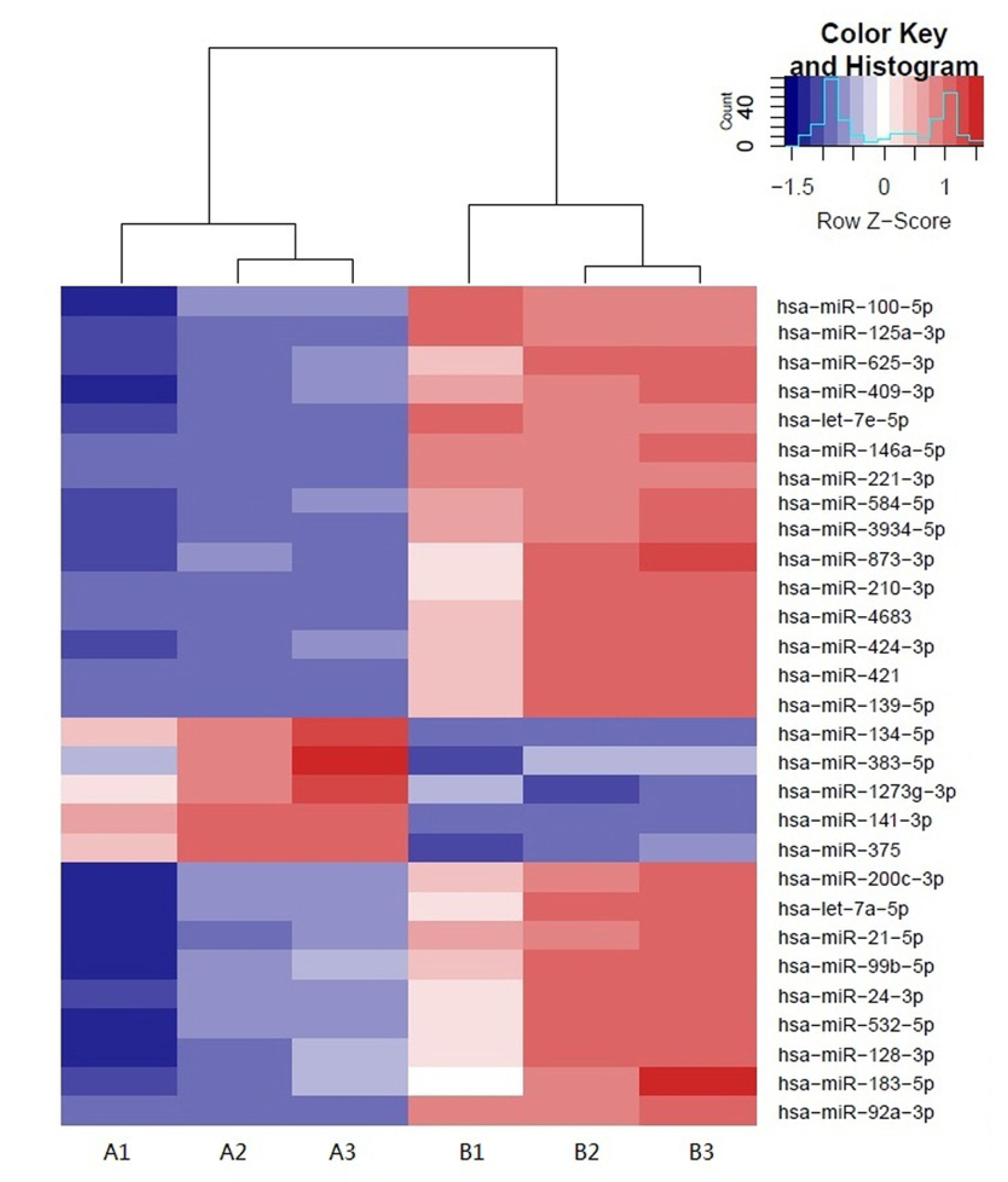
Heatmap of differences in miRNA expression levels.

**Fig 2.**
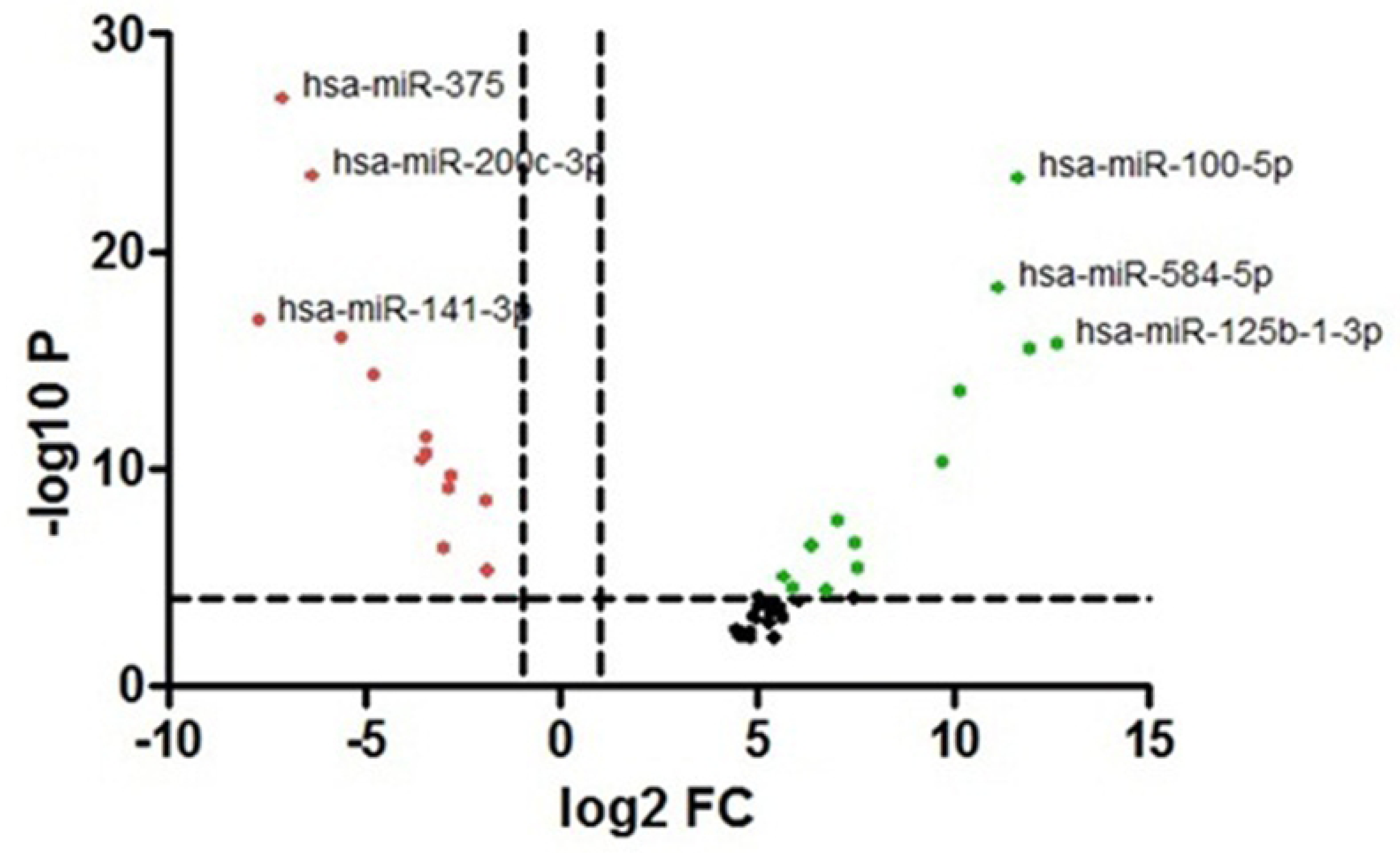
Volcano plot of differences in miRNA expression levels.

**Fig 3.**
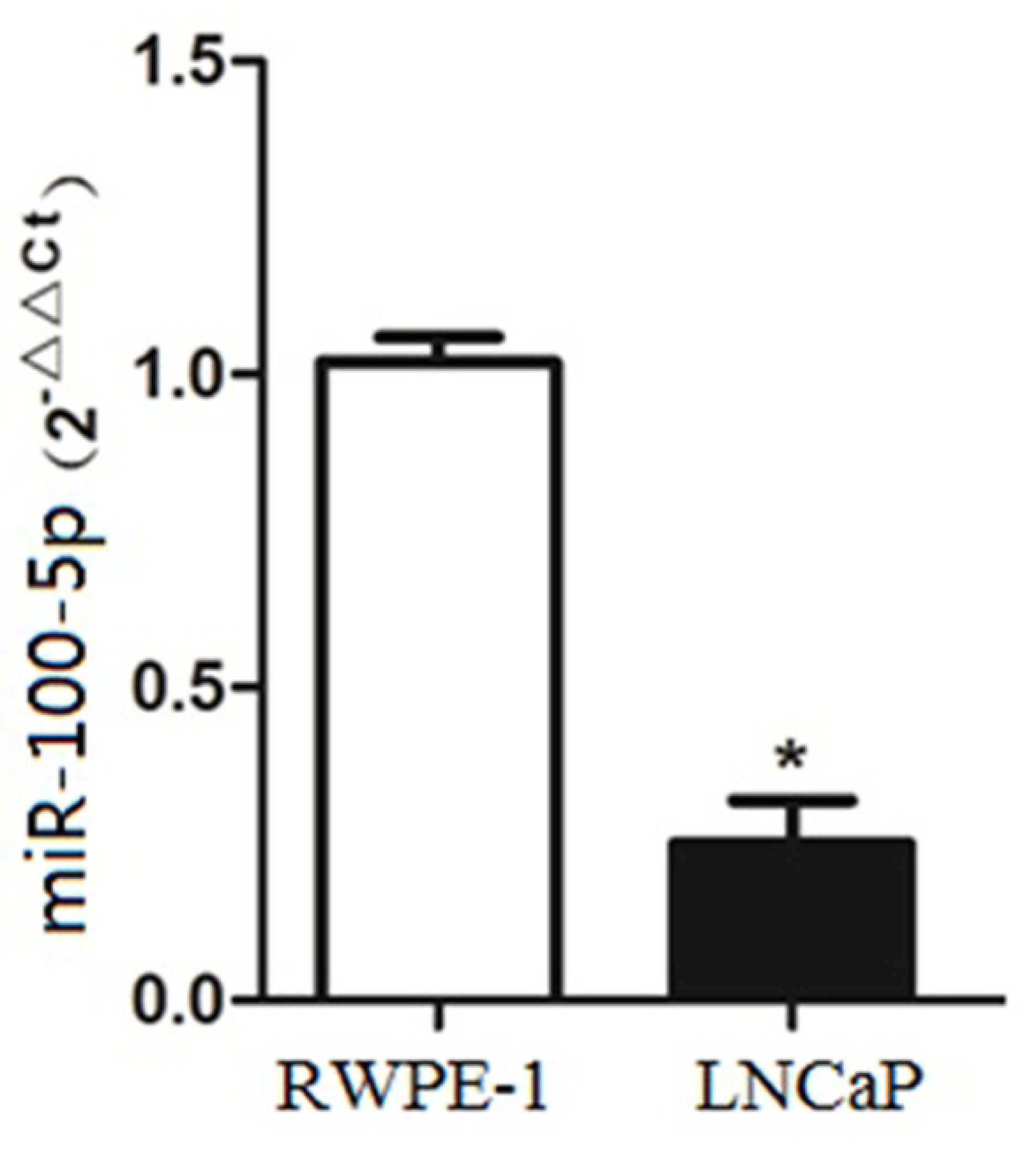
Expression of miR-100-5p in LNCaP and RWPE-1 cells. (* *P* < 0.01)

### Expression of miR-100-5p in LNCaP cells after transfection

The expression of miR-100-5p in the mimics group was significantly higher than that in the NC-mimics group (P<0.01) (figure 4), indicating that the transfection efficiency of miR-100-5p mimics was high.

**Fig 4.**
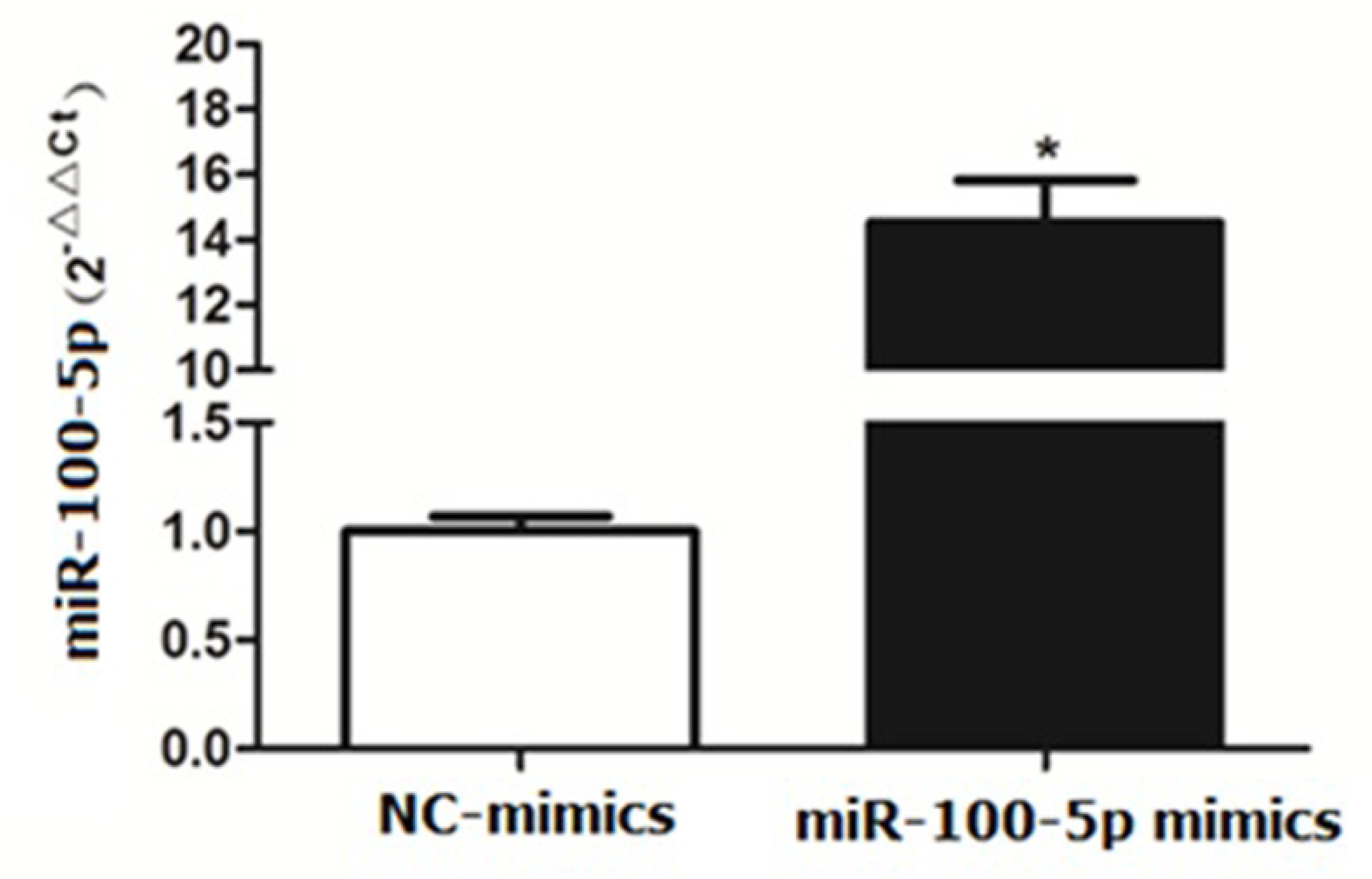
Expression level of miR-100-5p after transfection. (\**P* < 0.01)

### Proliferation activity of LNCaP cells after transfection

The CCK-8 proliferation assay showed that the OD values of LNCaP cells transfected with mimics at 48 h, 72 h and 96 h were significantly lower than those of the NC-mimics group (P<0.01) (figure 5).

**Fig 5.**
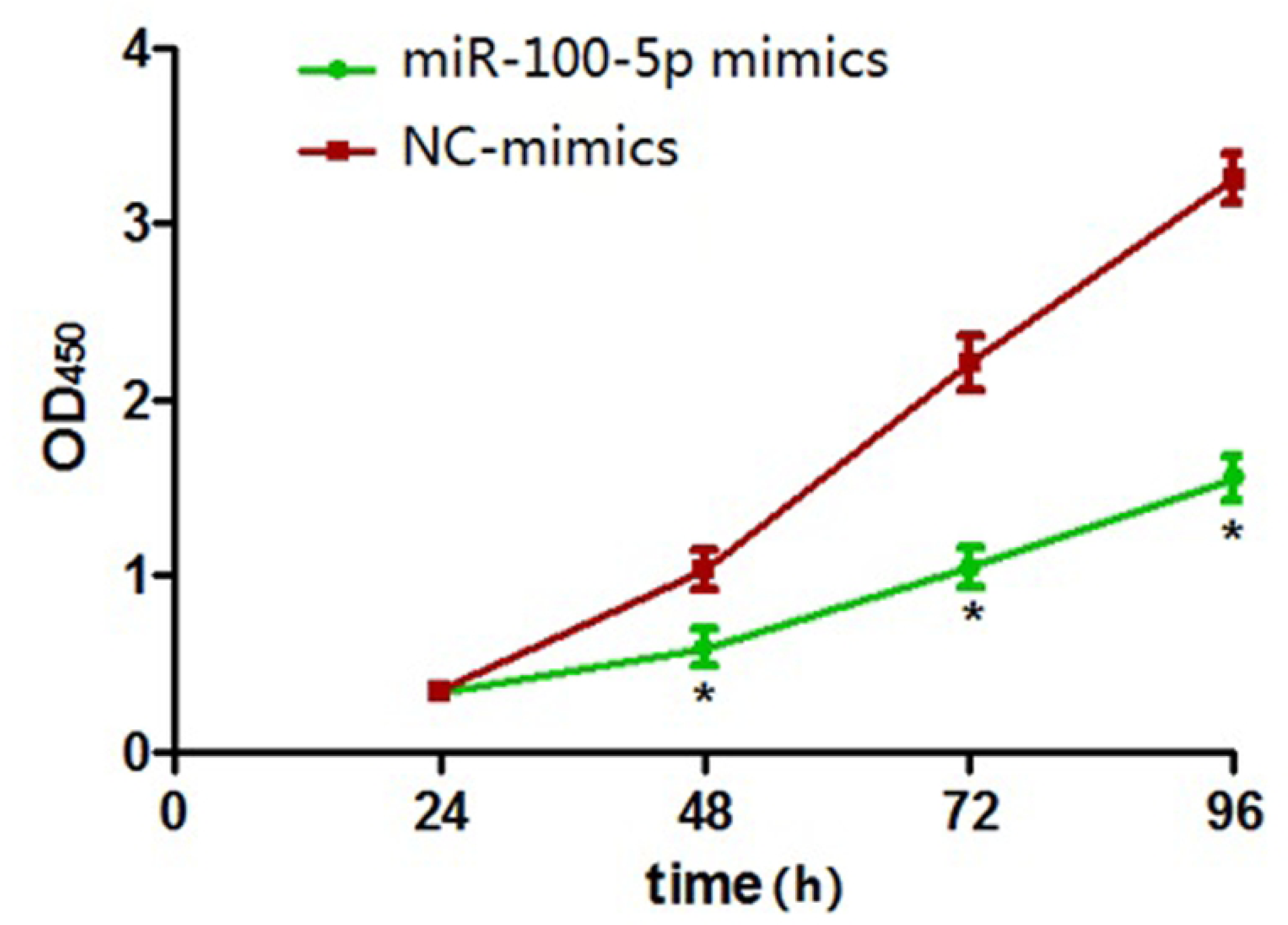
Cell proliferation activity after transfection. (\**P* < 0.01)

### Migration capacity of LNCaP cells after transfection

Compared with that of the NC-mimics group, the scratch healing rate of miR-100-5p mimics cells after transfection was significantly reduced (P<0.01), suggesting that the migration capacity of LNCaP cells after transfection was significantly reduced (figures 6) (figures 7).

**Fig 6.**
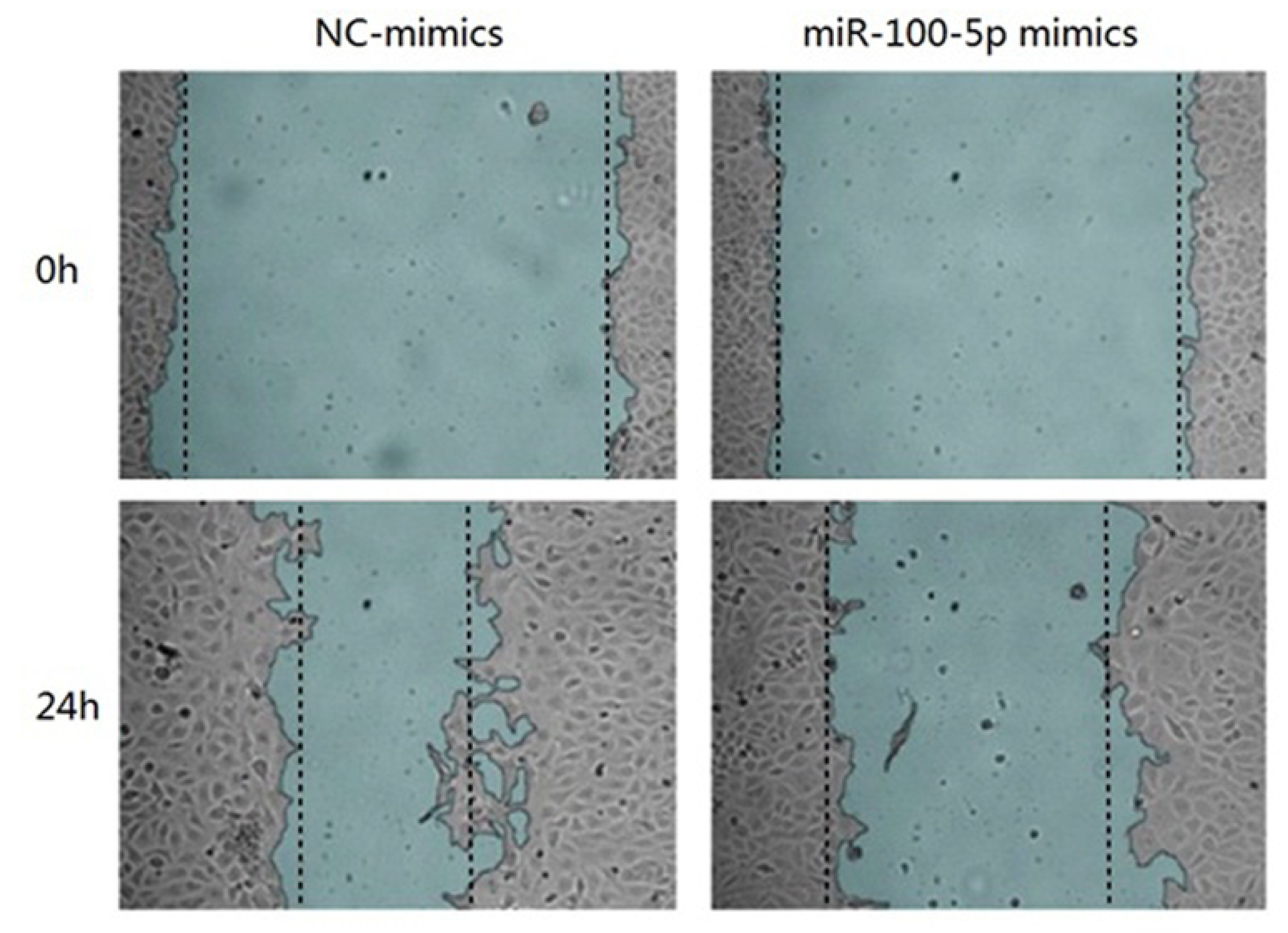
Cell migration ability detected by cell scratch experiment.

**Fig 7.**
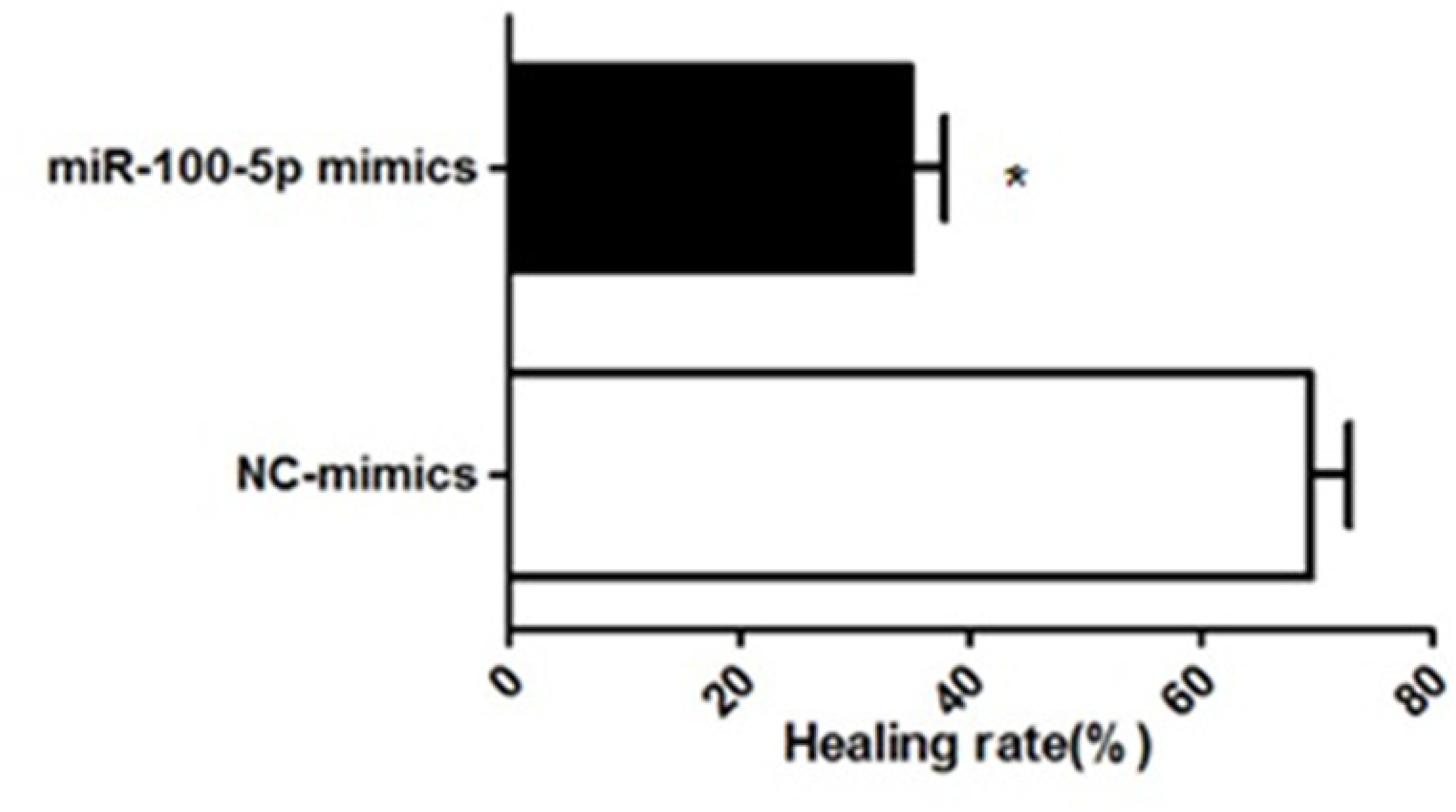
Change in the ability of cells to migrate.

### Invasion ability of LNCaP cells after transfection

The results of the Transwell assay showed that the number of membrane-penetrating cells in the miR-100-5p mimics group was significantly lower than that in the NC-mimics group (P<0.01), suggesting that the invasion capacity of transfected cells was significantly reduced (figure 8).

**Fig 8.**
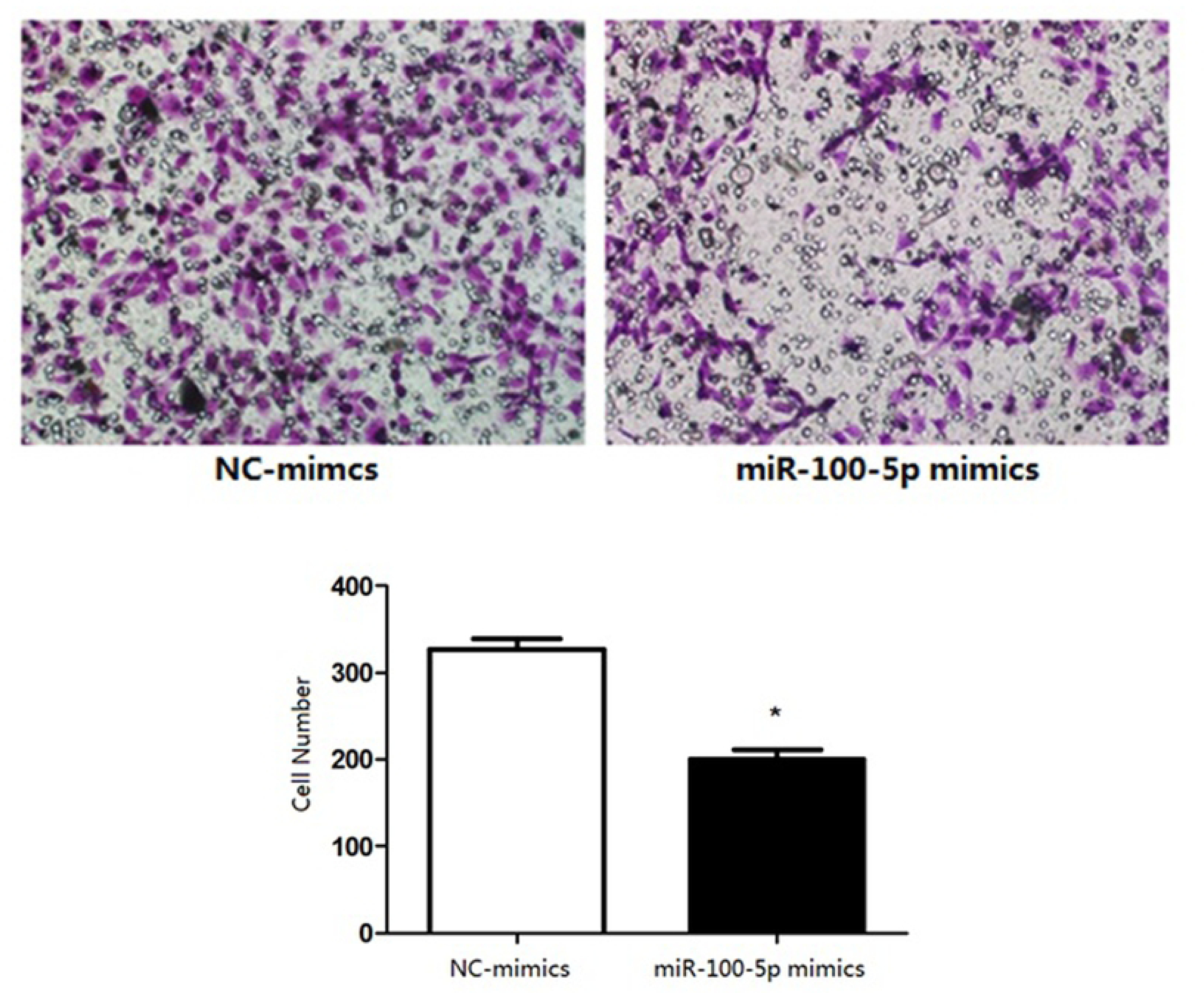
Invasion ability of cells detected by Transwell invasion assay. (\**P* < 0.01)

### Target gene prediction for miR-100-5p

A bioinformatics method was used to predict the potential target genes of miR-100-5p. By analysis in miRanda (http://www.microrna.org), mTOR was predicted to be a target gene of miR-100-5p, and there were complementary binding sites of seed sequences between miR-100-5p and the 3UTR of mTOR (figure 9).

**Fig 9.**
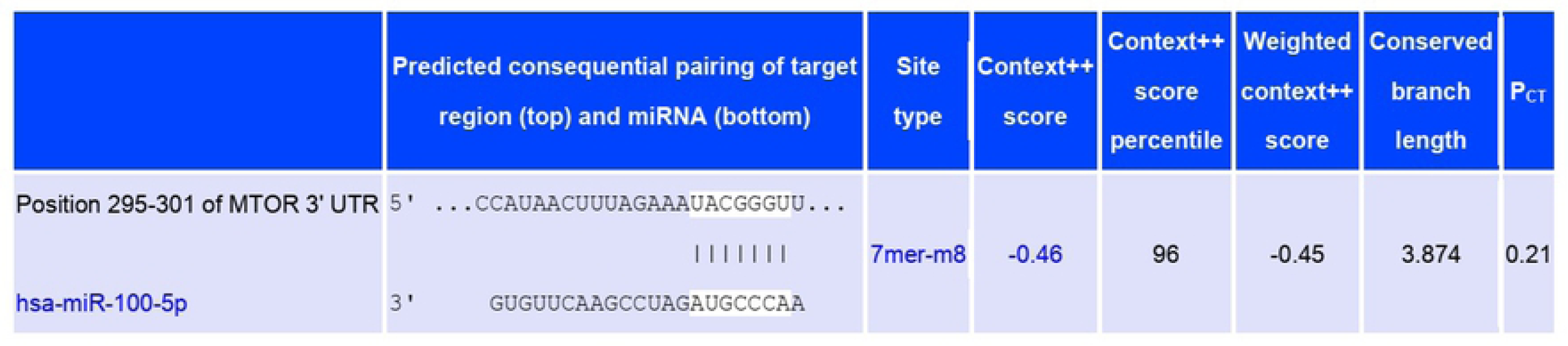
The binding sequence of miR-100-5p with the 3’UTR of mTOR.

### Expression of mTOR mRNA and protein

RT-PCR results showed that mTOR mRNA expression in LNCaP cells transfected with miR-100-5p mimics was significantly lower than that in the NC-mimics group (P<0.01). Western blot analysis showed that mTOR protein expression in LNCaP cells transfected with miR-100-5p mimics was significantly lower than that in the NC-mimics group (P<0.01), as shown in figure 10.

**Fig 10.**
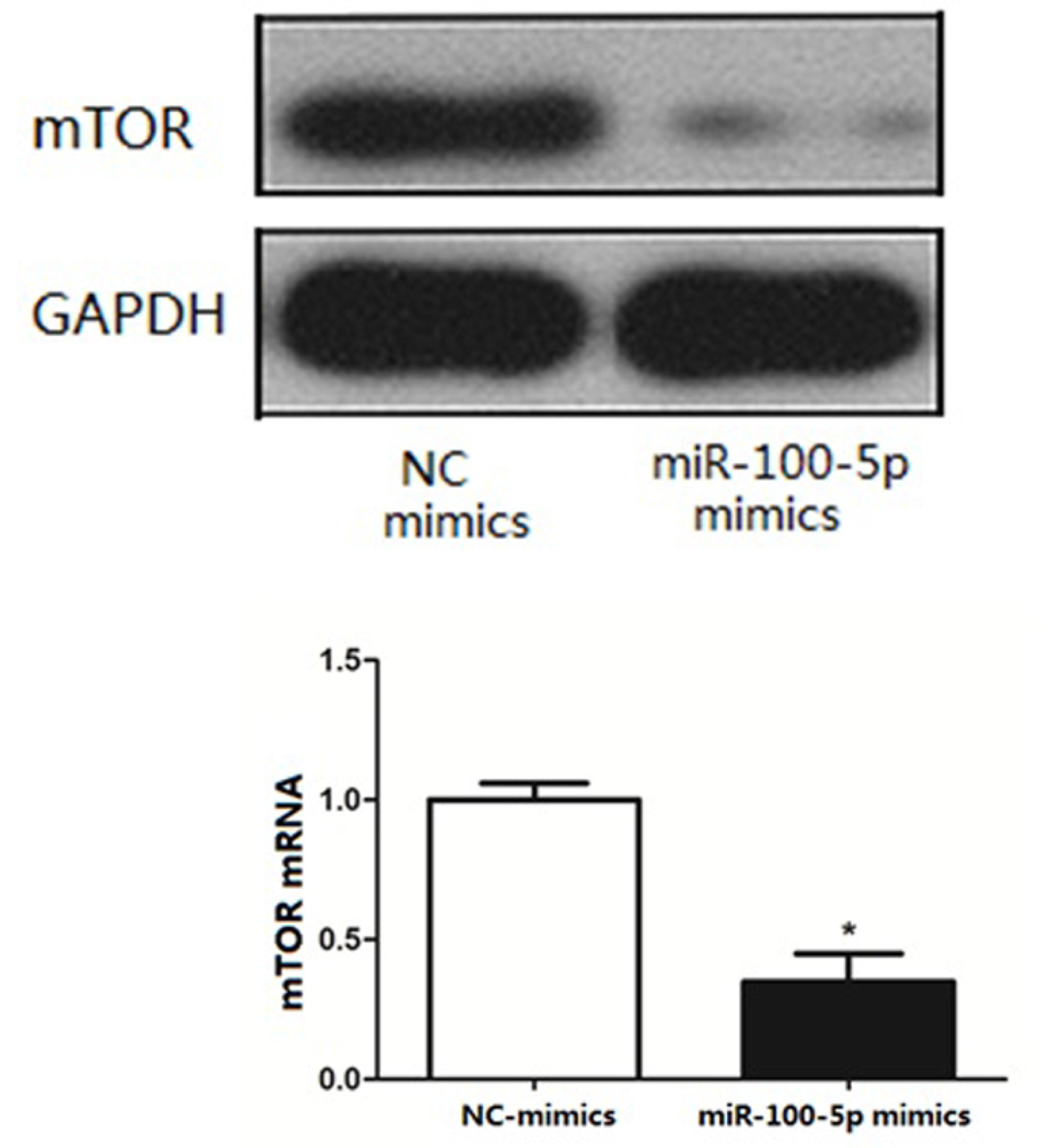
Expression of mTOR mRNA and protein after transfection. (\**P* < 0.01)

### Staining of bone tissue

Bone histopathological staining showed that miR-100-5p could significantly inhibit the occurrence and development of osteogenic bone metastases in prostate cancer (figure 11). Immunohistochemical results showed negative expression of mTOR in the bone tissue of group A and positive expression in the bone tissue of group B (figure 12).

**Fig 11.**
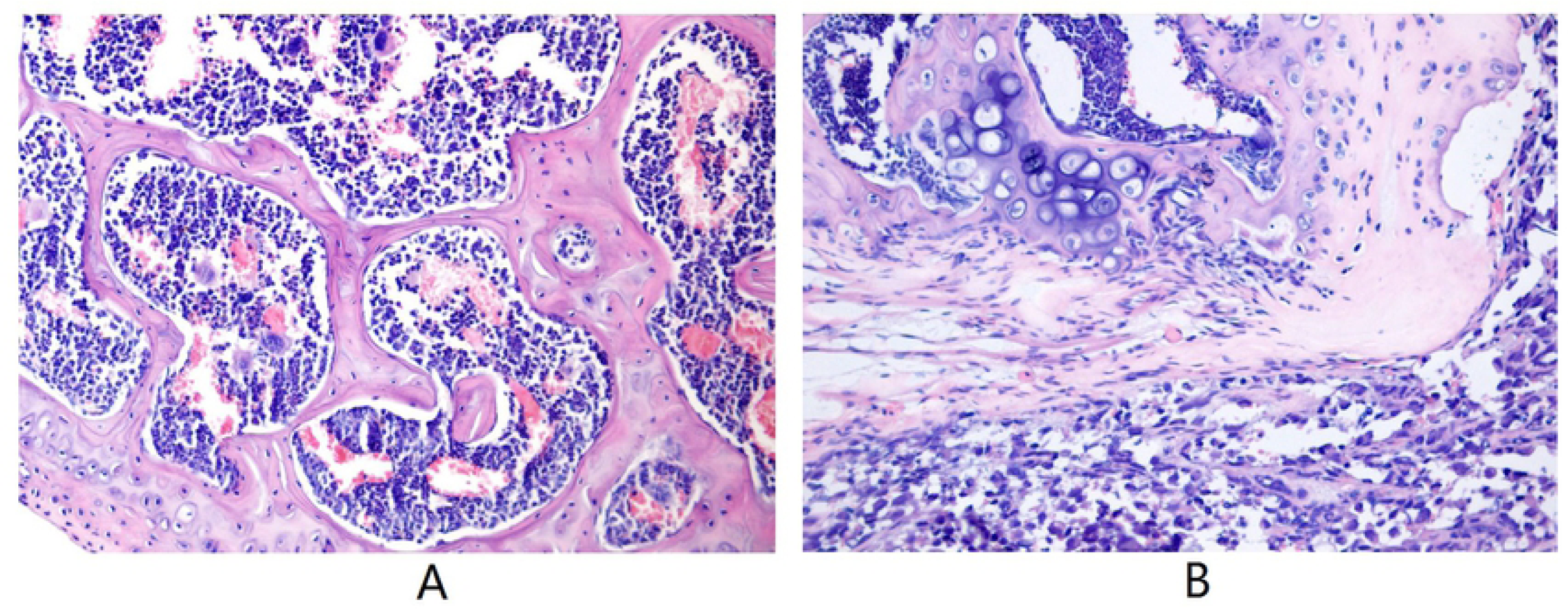
The result of HE staining.

**Fig 12.**
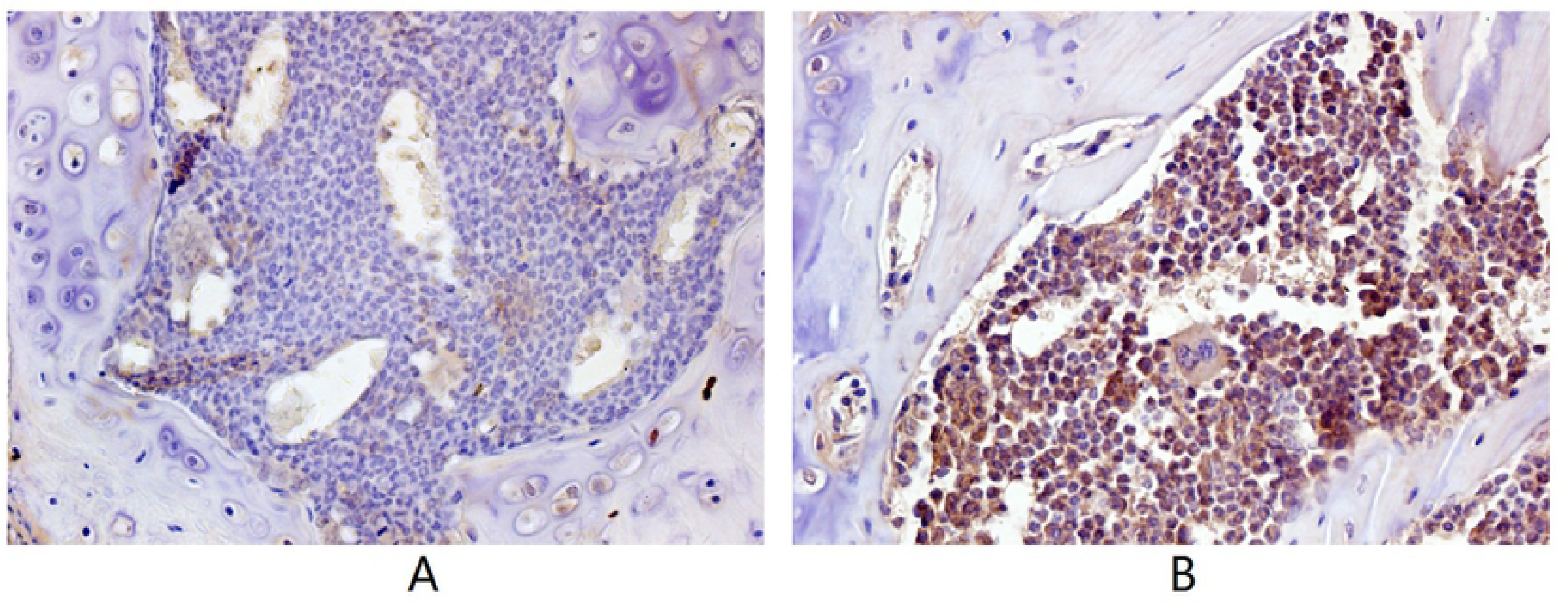
The results of immunohistochemical staining.

## Discussion

MiRNA was first discovered by Lee in 1993 and is a kind of ncRNA of approximately 19-24 nt.^11,12^ MiRNAs can bind to multiple target mRNA 3’UTRs and regulate gene expression, resulting in abnormal expression of target genes [13,14]. Abnormal miRNA expression has been associated with the occurrence and development of tumors. Studies have confirmed that differential expression of miRNAs in tumor tissues can be used as a biomarker for early detection, typing and prognosis of tumors.^15,16^ Studies have demonstrated altered levels of miRNAs in the development of PCa, with differences between PCa patients and healthy individuals.^17,18^ Studies have found that the expression of miR-100-5p is downregulated in prostate cancer, which is believed to be an important role of tumor suppressor genes in the development and progression of PCa.^19,20^ It has been reported that the absence of miR-100-5p leads to the upregulation of AGO2 expression levels, thus promoting the migration, invasion and EMT of cancer cells and promoting the metastasis of prostate cancer.^21^

In this study, the results of the NGS differential expression analysis showed that compared with those in RWPE-1 cells, miR-375, miR-200c-3p and miR-141-3p were upregulated, while miR-100-5p, miR-584-5p, and miR-125b-1-3p were downregulated. The results of the RT-PCR analysis further confirmed the low expression of miR-100-5p in LNCaP cells.

Some studies have found that miR-100-5p inhibits the proliferation, migration and invasion of tumor cells and the biological behaviors of tumors.^22,23^ To further study the effects of miR-100-5p on the proliferation, invasion and migration of LNCaP cells, we transfected miR-100-5p mimics into LNCaP cells by lipofection and successfully upregulated the expression of miR-100-5p in these cells. The results of the CCK-8 proliferation assay, cell scratch assay and Transwell invasion assay showed that the upregulation of miR-100-5p could significantly inhibit the proliferation activity, migration and invasion of tumor cells, further confirming that miR-100-5p, as a tumor suppressor gene, could inhibit the progression of PCa.

The downstream molecular pathway by which miR-100-5p inhibits the proliferation, invasion and migration of tumor cells is not completely clear. In studying the downstream target genes regulated by miR-100-5p in LNCaP cells, we found that the 3’UTR of miR-100-5p and mTOR has complementary seed sequence binding sites by using a bioinformatics method. Therefore, we predicted that mTOR might be a functional target gene of miR-100-5p. MTOR is a serine/threonine protein kinase that participates in physiological processes and pathological reactions by regulating protein synthesis and is related to the pathogenesis of cancer.^24^ The MTOR signaling pathway mainly regulates cell proliferation and metabolism involved in tumor development and is an important signaling pathway related to human cancer.^25^ In this study, we found that the expression levels of mTOR mRNA and protein decreased obviously in LNCaP cells after upregulation of miR-100-5p expression.

Next, nude mice in the two groups were administered by intraosseous injection with tumor cells. Group A cells were transfected with miR-100-5p mimics, and group B contained untransfected LNCaP cells (control). Pathological staining and immunohistochemical staining were performed after two months. HE staining of bone in group B showed a large amount of osteogenesis with tumor characteristics in the limbic cortex. The tumor cells were closely aligned, with loss of polarity and an imbalance in the nuclear to cytoplasmic ratio, and there was increased pathological nuclear fission. In group A, the trabecular structure was complete, and an obvious bone marrow cavity structure was observed. Immunohistochemical results showed negative expression of mTOR in the bone tissue of group A and positive expression in the bone tissue of group B. These results further confirmed that upregulation of miR-100-5p expression could significantly inhibit the expression of mTOR and that miR-100-5p played a role in the occurrence and development of PCa through targeted regulation of mTOR.

## Conclusion

miR-100-5p was downregulated in PCa LNCaP cell lines, and miR-100-5p inhibited the proliferation, migration and invasion of LNCaP cells. The mechanism is related to the downregulation of mTOR gene expression, which may become a molecular target of PCa targeted therapy in the future.

## Acknowledgments

This study was supported by the Natural Science Basic Research Program of Shaanxi (Program No. 2020JM-607). The authors express their gratitude to the study participants and research personnel for their involvement in the study. The authors declare that there are no conflicts of competing interests regarding the publication of this article.

